# Phylogenomic Assessment of the Role of Hybridization and Introgression in Trait Evolution

**DOI:** 10.1101/2020.09.16.300343

**Authors:** Yaxuan Wang, Zhen Cao, Huw A. Ogilvie, Luay Nakhleh

## Abstract

Trait evolution among a set of species—a central theme in evolutionary biology—has long been understood and analyzed with respect to a species tree. However, the field of phylogenomics, which has been propelled by advances in sequencing technologies, has ushered in the era of species/gene tree incongruence and, consequently, a more nuanced understanding of trait evolution. For a trait whose states are incongruent with the branching patterns in the species tree, the same state could have arisen independently in different species (homoplasy) or followed the branching patterns of gene trees, incongruent with the species tree (hemiplasy). Another evolutionary process whose extent and significance are better revealed by phylogenomic studies is gene flow between different species. In this work, we present a phylogenomic method for assessing the role of hybridization and introgression in the evolution of polymorphic or monomorphic binary traits. We apply the method to simulated evolutionary scenarios to demonstrate the interplay between the parameters of the evolutionary history and the role of introgression in a binary trait’s evolution (which we call *xenoplasy*). Very importantly, we demonstrate, including on a biological data set, that inferring a species tree and using it for trait evolution analysis in the presence of gene flow could lead to misleading hypotheses about trait evolution.

## Introduction

Evolutionary biology began with the study of traits, and both descriptive and mechanistic explanations of trait evolution are key foci of macroevolutionary studies today. Trait evolution is often coupled with speciation, as in the case of Darwin’s finches, where the evolution of their beaks reflects adaptation to particular diets in an adaptive radiation [1–4]. Modern systematics synthesizes genomic data into informative species trees [5, 6], revealing the complex relationship between speciation and trait evolution. This is a welcome development as statistical methods for elucidating interspecific trait evolution without making use of the species tree can produce misleading results [7, 8].

Given a hypothesized species tree inferred from available data, trait patterns “congruent” with the tree may be parsimoniously explained as having a single origin in some ancestral taxon, and are shared by all descendants. However, many trait patterns are “incongruent” and may be examples of convergent evolution, where traits have been gained or lost independently. This kind of explanation is termed homoplasy, referring to a pattern of similarity which is not the result of common descent [9]. Incongruent trait patterns can also be produced by discordant gene trees and ancestral polymorphism. In such cases, while the trait pattern is incongruent with the species tree, it is congruent with gene trees that differ from the species tree.

When gene tree incongruence is due to incomplete lineage sorting (ILS) this explanation is termed hemiplasy [10, 11], and the hemiplasy risk factor (HRF) was developed to assess its significance for a given species tree [12]. Inference of species trees from genomic data in the presence of ILS has attracted much attention in recent years, resulting in a wide array of species tree inference methods [13–20]. The significance of elucidating not only the species tree but also the gene trees within its branches was recently highlighted for its significance in understanding trait evolution [21].

Another major source of species/gene tree discordance in eukaryotes is hybridization and introgression [22]. The multispecies network coalescent was developed to unify phylogenomic inference while accounting for both ILS and introgression [23–25]. Gene flow may explain some trait evolution [26], and methods analyzing trait evolution along a species network have been introduced [27, 28]. Such methods do not account for ILS, but the HRF framework was recently extended to fold introgression into hemiplasy and homoplasy [29]. However, hemiplasy was originally circumscribed to discordances that arise from idiosyncratic lineage sorting [11]. To distinguish the effects of gene flow we therefore propose using “xenoplasy” to explain a trait pattern resulting from inheritance across species boundaries through hybridization or introgression. This builds on “xenology” which denotes homologous genes sharing ancestry through horizontal gene transfer [30].

For the example in Fig 1, although both gene trees share the same topology, mutations along the internal branches will lead to hemiplasy or xenoplasy respectively for the solid and dashed gene trees. It also illustrates that hemiplasy requires deep coalescence events, but xenoplasy does not. It is important to highlight here that in some cases there cannot be clear delineation of homoplasy, hemiplasy, and xenoplasy, as the evolution of trait could simultaneously involved convergence and genes whose evolutionary histories involve both ILS and introgression. In fact, the picture can get even more complex when the effects of gene duplication and loss are involved (maybe necessitating yet another term, e.g., “paraplasy,” following the term “paralogy” that is used to describe genes whose ancestor is a duplication event).

Figure 1: Phylogenomic view of trait evolution in the presence of incomplete lineage sorting (ILS) and introgression. Left: The three possible genealogies of three taxa A, B, and C. Right: Phylogenetic network that models an underlying species tree (A,(B,C)) along with a reticulation from A to B, and whose associate inheritance probability is *γ*. The embedded solid gene tree involves ILS but no introgression, whereas the dashed gene tree involves introgression but not ILS. The states *S_a_*, *S_b_*, and *S_c_* of an incongruent binary character are shown at the leaves of the phylogenetic network.

We introduce the global xenoplasy risk factor (G-XRF) to assess the role of introgression in the evolution of a given binary trait. We append “global” because unlike HRF, which is computed per-branch, G-XRF is computed over the whole network for a specific pattern, a pattern which can be polymorphic. We evaluated the G-XRF in simulated settings with ILS and introgression, demonstrating the interplay among divergence and reticulation times, introgression probability, population size and substitution rates, and how this affects the role of introgression in trait evolution. We also show how sampling trait polymorphism improves the informativeness of the G-XRF, and the importance of inferring a species *network* where gene flow occurs for elucidating trait evolution. In particular, we demonstrate how assuming a species *tree* despite the presence of gene flow overemphasizes the role of hemiplasy.

Our work brings together phylogenetic inference and comparative methods in a phylogenomic context where both the species phylogeny and the phylogenies of individual loci are all taken into account. A short tutorial demonstrating how to calculate and use G-XRF values is available at our web site, https://nakhlehlab.github.io/.

## Materials and methods

### The Global Xenoplasy Risk Factor

Consider that a binary trait evolving along the branches of a fixed species tree or network Ψ with population mutation rates Θ, and in the case of species networks inheritance probabilities Γ. The trait is given by 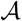 which specifies for each species the number of sampled individuals with state 0 and the number with state 1. We refer to this as the **observed state counts**, or in the special case where only one observation present for each species, as the **trait pattern**. We use *u* and *v* respectively for the forward character substitution rate (replacing state 0 with state 1) and the backward character substitution rate (replacing state 1 with state 0).

The posterior probability of the species phylogeny and associated parameters given 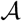 is:

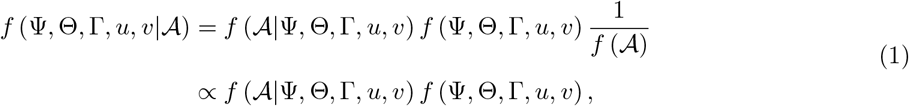

where 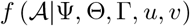 is the likelihood of the observed state counts, and *f* (Ψ, Θ, Γ*, u, v*) is the prior on the species phylogeny and population sizes.

In the phylogenomic view of trait evolution, the evolutionary history of 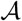 is modeled as a gene tree evolving inside the species phylogeny. To calculate the likelihood of the observed state counts, we need to integrate over all possible genealogies *G*:

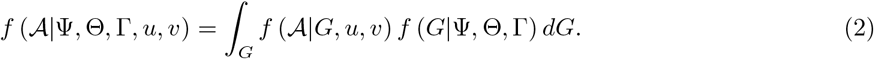

Here, 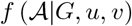 is the likelihood of a genealogy given the observed site counts and *f* (*G|*Ψ, Θ, Γ) is the multispecies coalescent (or multispecies network coalescent) likelihood. We use existing Bayesian methods of species tree and network inference from bi-allelic markers [31, 32] to calculate 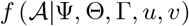 according to Equation 1. While the network inference method we use cannot handle missing data, it can calculate the likelihood where multiple individuals are sampled for a single species, which we take advantage of to calculate the likelihood of polymorphic traits. Finally, the G-XRF is calculated as the natural log of the posterior odds ratio, where Ψ is the species network which should be estimated from the data, and 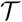 is the hypothesized backbone tree without gene flow displayed by Ψ:

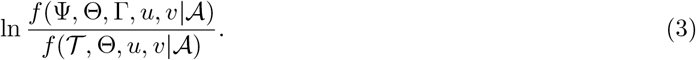

This ratio compares the posterior probability integrating over possible hemiplasy, homoplasy and introgression with the probability integrating over possible hemiplasy and homoplasy alone. Therefore, the ratio compares how likely it is that introgression has contributed to the trait pattern, rather than directly comparing introgression with hemiplasy or introgression with homoplasy.

### Jaltomata analysis

We studied the utility of G-XRF by inferring species phylogenies from a previously published dataset of 6,431 orthologous gene sequences from *Jaltomata* and the close relative *Solanum lycopersicum* as an outgroup [33]. To derive conditionally independent bi-allelic markers of the original dataset, we randomly selected one site from each gene and obtained 6,409 valid bi-allelic markers in total.

We inferred a species phylogeny of this group in two different ways using MCMC BiMarkers [32] with chain length 5 *×* 10^6^, burn-in 2 *×* 10^6^, and sample frequencies 1000, using the following command:

~~~
MCMC_BiMarkers -taxa (JA0701, JA0456, JA0694, JA0010, JA0719, JA0816)
-cl 5000000 -bl 2000000 -sf 1000 -mr 1
~~~

We ran the same command setting `-mr` to 0 (which sets the number of reticulations to 0) for species tree inference. The *effective sample size* (ESS) of the parameter values of the MCMC chains were higher than 2321 for the species tree and higher than 1583 for the species network.

### Simulated multilocus data

We generated the data with 2 steps. First, we generated 128 gene trees with ms [34] given the species network. The command is as follows.

~~~
ms 6 128 -T -I 6 1 1 1 1 1 1 -es 0.25 5 0.3 -es 0.25 3 0.8 -ej 0.5 7 3
-ej 0.5 8 2 -ej 0.75 6 5 -ej 1.0 3 4 -ej 1.0 2 1 -ej 2.0 5 4 -ej 2.5 4 1
~~~

Second, at each locus, we simulated the sequence alignment given the gene tree with seq-gen [35]. We set the length of sequences to be 500 bps, and utilized GTR model with base frequencies 0.2112,0.2888,0.2896,0.2104 (A,C,G,T) and transition probabilities 0.2173,0.9798,0.2575,0.1038,1.0,0.207. We set the population mutation rate *θ* = 0.036, so the scale *−s* is 0.018. The command is as follows.

~~~
seq-gen -mGTR -s0.018 -f0.2112,0.2888,0.2896,0.2104
-r0.2173,0.9798,0.2575,0.1038,1.0,0.207 −l500
~~~

We inferred a species network from the simulated data with MCMC SEQ [36] under GTR model with chain length 5 *×* 10^7^, burn-in 1 *×* 10^7^ and sample frequencies 5000. We fixed the population mutation rate *θ* = 0.036 and GTR parameters to be true parameters. The command is below:

~~~
MCMC_SEQ -cl 60000000 -bl 10000000 -sf 5000 -pl 8
-tm <A:A_0;C:C_0;G:G_0;L:L_0;Q:Q_0;R:R_0> -fixps 0.036
-gtr (0.2112,0.2888,0.2896,0.2104,0.2173,0.9798,0.2575,0.1038,1,0.2070);
~~~

We also inferred a species tree using StarBEAST2 [17]. The chain length was 10^8^ with a sample frequency of sample frequency 50, 000 under GTR model with empirical base frequencies and transition probabilities fixed to the true values. Population sizes were sampled for the individual branches (i.e., a single population size across all branches was *not* assumed).

## Results

Consider the evolutionary history depicted by the phylogenetic network of Fig 1. If a single individual is sampled from each of the three species A, B, and C, then this network can be viewed as a mixture of two displayed trees [37]: The “species” tree (A,(B,C)) and another tree that captures the introgressed parts of B’s genome ((A,B),C). The given trait whose character states are 1, 1, and 0 for taxa A, B, and C, respectively, could have evolved down and within the branches of the species tree. In this case, either homoplasy and hemiplasy could explain the trait evolution. To tease these two processes apart, assuming introgression did not play a role, the HRF can be evaluated with respect to the species tree. Furthermore, a similar analysis of both displayed trees can provide a way for assessing the role of hemiplasy in the presence of introgression [29]. In our case, we are interested in answering a different question: How much does a reticulate evolutionary history involving hybridization and introgression explain the evolution of a trait as opposed to a strictly treelike evolutionary history?

The likelihood of observed state counts given the species phylogeny integrates over all possible gene histories and is calculated using methods previously implemented in PhyloNet [32, 38]. Furthermore, while the model was illustrated above on three taxa, those methods allow for any number of taxa and any topology of the phylogenies, including any number of reticulation events. We use G-XRF to measure the importance of taking into account the possibility of introgression for a given trait. The higher value of G-XRF corresponds to the greater necessity of a species network for trait analysis, and the greater odds that the site pattern is due to introgression.

### Interactions between evolutionary parameters

A phylogenomic view of the evolution of a binary trait on the phylogenetic network of Fig 1 involves, in addition to the topologies of the phylogenetic network and species tree, roles for:

- The inheritance probability *γ*, which measures the probability that a locus in the genome of B was derived from the ancestor of A, representing gene flow from A into B [24, 36].
- The reticulation time *T_r_*, as it controls the likelihood of inheriting a character state by B from A, as well as the likelihood of such an inherited state becoming fixed in the population.
- The length of the internal species tree branch, *T*_2_ *− T*_1_, as it controls the amount of ILS and, consequently, hemiplasy.
- The population mutation rate, *θ* = 2*N*_2_*μ*, which also controls the amount of ILS and hemiplasy.
- The relative forward and backward substitution rates *u, v*.

The character states are shown at the leaves of the network of Fig 1 which displays the species tree (A,(B,C)). We varied the ILS level by varying the internal branch length (*T*_2_ *− T*_1_). The initial interval between internal nodes *T_n_* was 1 coalescent unit, but we varied (*T*_2_ *− T*_1_) from 0.001 to 10 to represent a range from very high to very low levels of ILS. Two factors controlled the introgression: the inheritance probability *γ* and the reticulation time *T_r_*. The inheritance probability *γ* was varied between 0 and 1. As *γ* approaches 1 this represents a complete replacement of the genome with introgressed sequences, as seen in the *Anopheles gambiae* species complex [39]. The reticulation time *T_r_* was varied between 0 and 1 coalescent unit. We varied the population mutation rate *θ* between 0.001 and 0.01. For the character substitution rate, we used three settings: forward = 0.1*×*backward, forward = backward and forward = 10*×*backward. For the polymorphic trait, we varied the frequency of allele ‘1’ in taxon B from 0 to 1.

We focused on a couple of three-way interactions: G-XRF as a function of the interplay among the internal branch length, the inheritance probability, and the relative forward/backward character substitution rates (Fig 2 top row), and G-XRF as a function of the interplay among the reticulation time, population mutation rate, and the relative forward/backward character substitution rates (Fig 2 bottom row).

**Figure 2:**
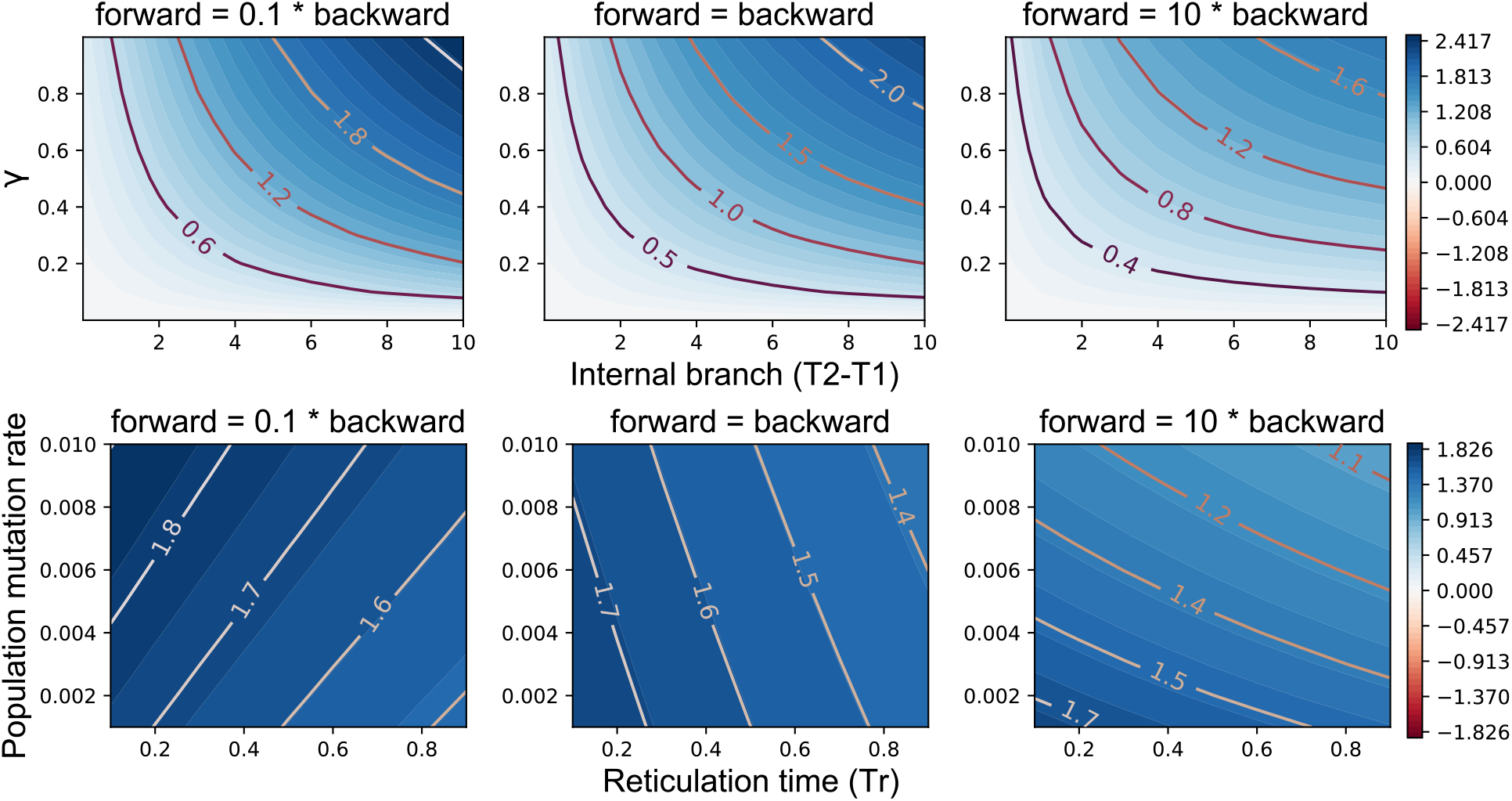
The interaction of evolutionary parameters affects the need for introgression to explain trait patterns. G-XRF is shown as a function of internal branch length *T*_2_ − *T*_1_ and inheritance probability *γ* when reticulation time *T_r_* = 0:1 coalescent units and population mutation rate *θ* = 0:01 (top row), and as a function of *θ* and *T_r_* when *T*_2_ − *T*_1_ = 10 and *γ* = 0:5 (bottom row).

As the internal branch becomes longer, the amount of ILS and consequently hemiplasy decrease, increasing the roles of introgression/homoplasy. Conversely, as the forward substitution rate increases relative to the backward rate, the necessity of introgression decreases since convergent mutations along the A and B branches may explain the trait pattern. This is indicated by decreasing G-XRF values for the same combination of (*T*_2_ − *T*_1_) and *γ* across as forward substitution rate increases (Fig 2 top row).

The second three-way interaction is based on a scenario where the internal branch is too long for ILS to occur and, consequently, for hemiplasy to be a factor. Therefore, the two forces underlying trait evolution in this case are homoplasy and xenoplasy. The role of introgression increases as *T_r_* decreases, since there is less time for the state to revert to 0 when state 1 is inherited by B from its most recent common ancestor (MRCA) with A (Fig 2 bottom row). The other key factor is the probability of a forward mutation, which is a function of the population mutation rate and the ratio of forward to backwards mutations. As this probability increases, homoplasy becomes more plausible as an explanation through convergent forward mutations along the A and B branches the same as for the first three-way interaction.

Increasing the probability of forward relative to backwards mutation flips the effect of increasing the population mutation rate *θ*. When the probability of forward mutation is low (and backward mutation high), increasing *θ* makes the trait pattern more likely to be the result of introgression, since any mutations along the B branch are likely to be backward (Fig 2 bottom left). When the probability of forward mutation is high (and backward mutation low), increasing the population mutation rate makes homoplasy more plausible due to convergent forward mutations along the A and B branches (Fig 2 bottom right).

### Introgression and polymorphic traits

Polymorphism is a major factor in trait evolution, often ignored only because methods do not account for it [40]. Fortunately, bi-allelic marker methods based on the multispecies (network) coalescent methods naturally account for polymorphism, and we take advantage of that in order to apply G-XRF to polymorphic traits. We conducted the same analysis as above, but now with ten observations for taxon B (we assume only one sampled state each from taxa A and C). Once again the internal branch is too long for ILS and hemiplasy to be relevant to the results.

Under certain conditions the G-XRF values were much higher or lower than what we observed sampling only one state per species (Figs 3 and 4). This is predictable, as we now have 12 total observations of the trait state compared with only three observations before, and more data will increase the magnitude of the observed state count likelihoods.

**Figure 3:**
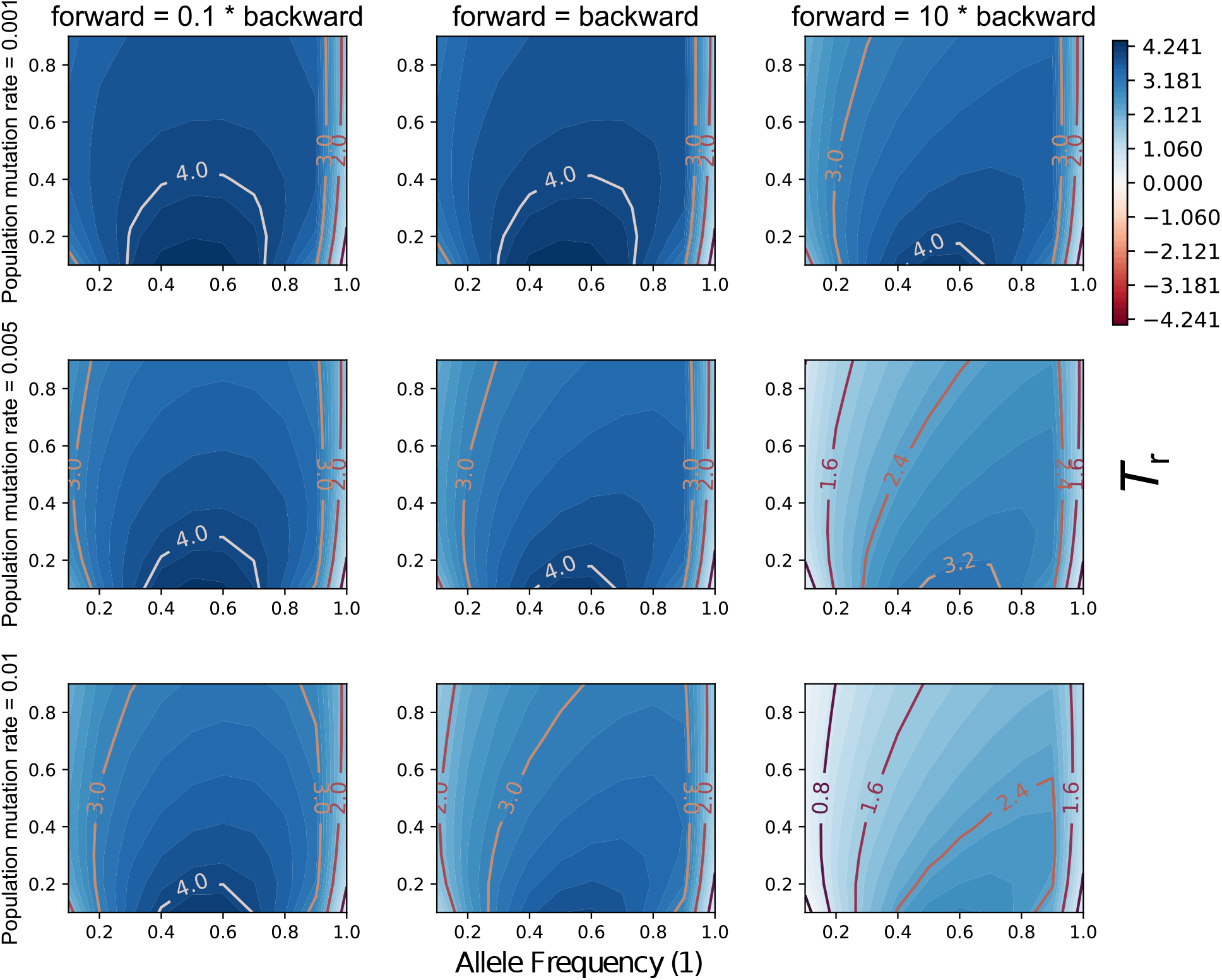
The interaction of evolutionary parameters affects the need for introgression to explain observed state counts. The x- and y-axis in each panel correspond to the frequency of character state 1 in taxon B and the reticulation time *T_r_*. Columns correspond to three different relative forward/backward character substitution rates and rows correspond to three different population mutation rates. In all panels *T*_2_*−T*_1_ = 10 coalescent units and *γ* = 0.5.

**Figure 4:**
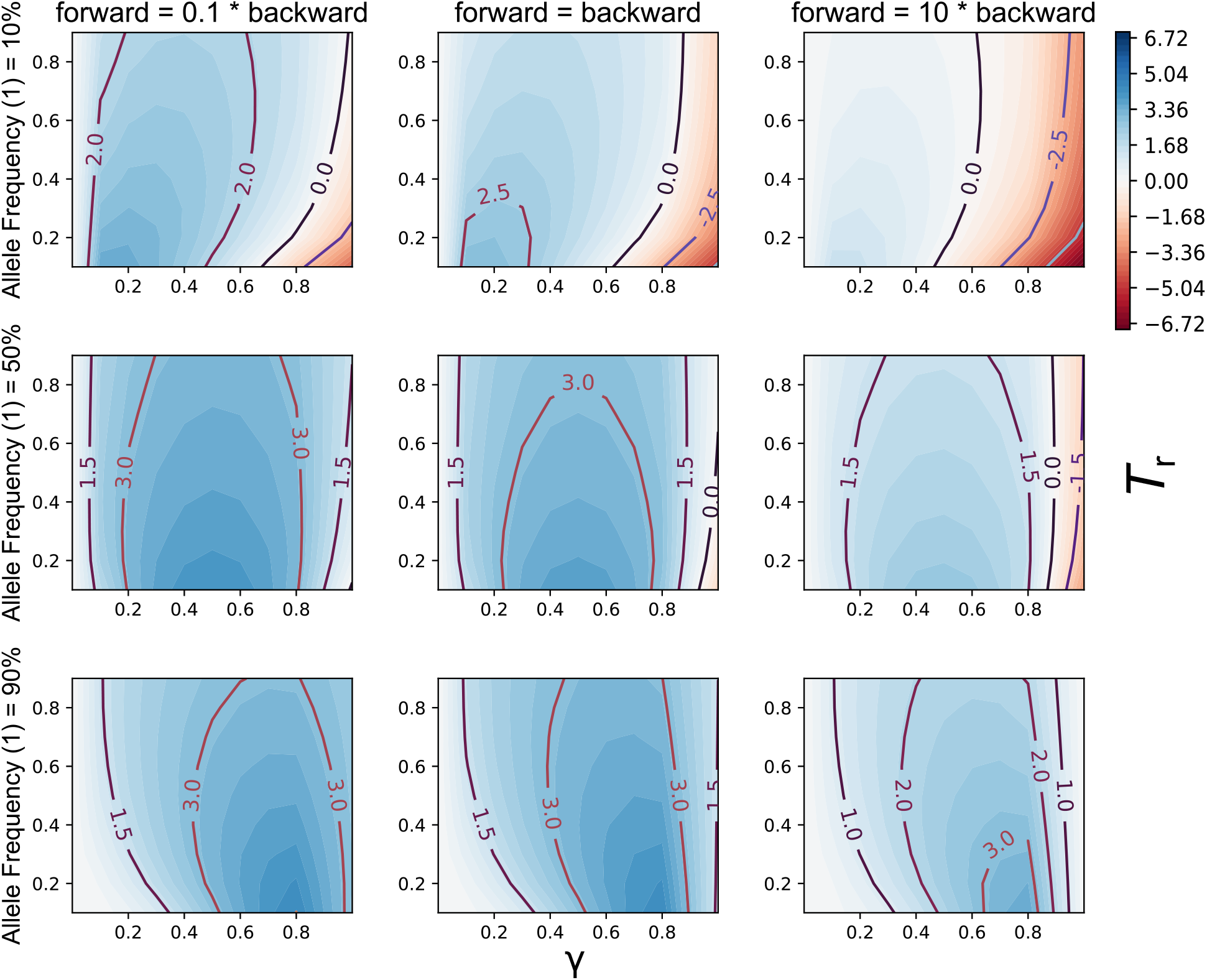
G-XRF values in the presence of trait polymorphism. The x- and y-axis in each panel correspond to the inheritance probability *γ* and reticulation time *T_r_*, respectively. Columns correspond to three different relative forward/backward character substitution rates, and rows correspond to three different frequencies of all 1 in taxon B. In all panels *T*_2_ *− T*_1_ = 10 coalescent units and *θ* = 0.01.

The G-XRF is highest where the introgression probability *γ* is equal to the observed frequency of the 1 state in B, an intuitively predictable result (Fig 4). Increased population mutation rate decreased the G-XRF, especially when the forward substitution rate was relatively high and the frequency of 1 in B relatively low (Fig 3). As for the previous results, this is because convergent forward mutations may occur along the A and B branches. Unlike for trait patterns with only one observation per species, we can now observe negative G-XRF values. When the observed frequency of 1 in B is low, but *γ* is high, the trait is much more plausibly explained through common ancestry between B and C than gene flow (Fig 3). This effect becomes stronger as the probability of forward mutation increases, as it makes backward mutation of introgresses traits less likely.

### Applying G-XRF to *Jaltomata*

When the evolutionary history of a set of species is reticulate, inferring a species tree could result in a tree with much shorter branches [25, 36, 41]. In such cases, the role of hemiplasy would be overestimated as it has an inverse relationship to branch length. This could in turn give the false impression that introgression did not play a role in the trait’s evolutionary history. In other words, inferring a species tree despite the presence of gene flow could lead to misleading results not only in terms of the evolutionary history of those species, but also for their associated traits.

We illustrate this phenomenon using empirical and simulated data. Based on an inferred species tree, the trait patterns of *Jaltomata* species were previously hypothesized to be the result of homoplasy [42]. Another study indicated that the evolutionary history of these species was reticulate, yet no phylogenetic network was inferred [33]. We inferred both a species tree and species network based on six *Jaltomata* species and the *Solanum lycopersicum* outgroup from the latter study (Fig 5).

Figure 5: Inferred species tree (left) and network (right) of the *Jaltomata* data set. The major tree inside the species network is obtained by removing the blue reticulation edge leading to I1.

We evaluated the HRF values of the species tree inferred without reticulations, and of the major tree inside the species network. The HRF values computed based on the species tree are larger than the values computed based on the major tree inside the species network. This suggests that the predicted amount of hemiplasy is erroneously high when gene flow is unaccounted for. We also computed G-XRF for three possible trait patterns, finding that trait patterns X and Y can be plausibly explained by either tree-like or reticulate evolution since the G-XRF values are close to zero (Fig 6). The trait pattern that would be best explained by introgression was pattern Z where introgression of state 1 from the MRCA of (*incahuasina*, *grandibaccata*, *dendroidea*) into the MRCA of (*procumbens*, *repandidentata*) would be a more plausible explanation than homoplasy, except for when the probability of forward mutation is relatively high and therefore convergent forward mutations can be anticipated.

**Figure 6:**
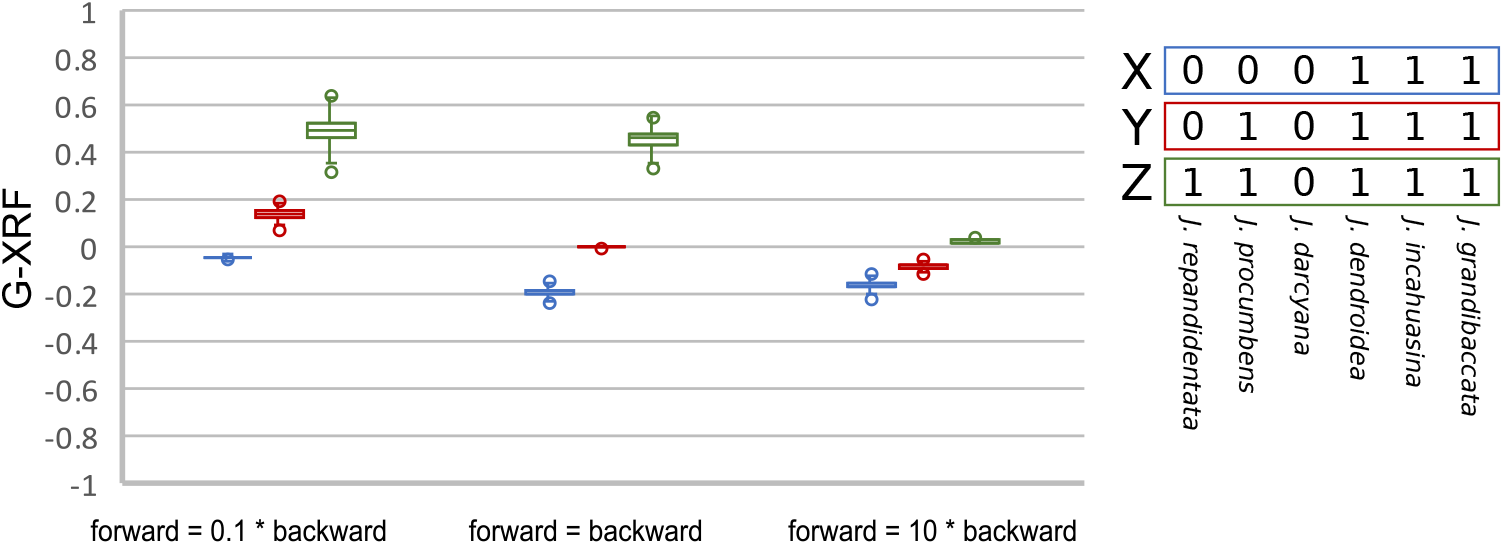
G-XRF values of three trait patterns (X, Y and Z) as the ratio of forward to backward substitutions is varied. Each box plot summarizes 3,000 G-XRF values obtained from the species network and corresponding major tree sampled from the posterior distribution of *Jaltomata* species networks.

### The simulated data set

To further confirm these results, we repeated the same analysis on simulated data. We simulated sequence alignments on 128 loci from the phylogenetic network whose topology was based on a previously published phylogeny of anopheline mosquitoes [43]. Then, we inferred a species network and tree from the simulated alignments. We then computed HRF values on two trees:

- The “major tree” of the species network estimated by, obtained by deleting the edge with the lowest inheritance probability entering each reticulation node. Specifically, the reticulation edges *I*5 *→ I*6 and *I*7 *→ I*8 were deleted as they have the smaller inheritance probabilities.
- The inferred species tree. Unlike the major tree, this was not ultrametric in coalescent units, because we did not assume a single uniform population size across all branches in this case.

The major tree HRF values for the branches leading to the two clades of three *Jaltomata* species each were orders of magnitude smaller than the HRF values for the same branches in the species tree. This indicates that some of the gene tree incongruence is erroneously attributed to ILS, and that incongruent trait patterns may erroneously be attributed to hemiplasy, when introgression is not accounted for.

We also compare posterior probability densities for the case where taxa A and C have state ‘1’ and the other taxa have states ‘0’ and the case where Q and R have state ‘1’ and the other taxa have states ‘0’. Both cases are examples of where introgression from the second taxon’s lineage to the first taxon’s lineage could explain the trait pattern. We find that the probability density of the major tree is lower than the true or inferred networks in either case, suggesting that the G-XRF is powerful enough to detect the potential for specific traits to be introgressed, since it is derived from those probability densities. Similar posterior probabilities for the true and inferred networks further suggest that relying on inferred species phylogenies to compute the G-XRF is not a problem.

## Discussion

The extent of hybridization and introgression continues to be revealed in an increasingly larger number of eukaryotic clades [44]. In this paper, we introduced the concept of xenoplasy to capture the inheritance of morphological character states via hybridization and introgression. We demonstrated how various evolutionary parameters impact the role these processes could play in the evolution of a given trait, including polymorphic traits. When gene flow is ignored as a mode of inheritance, complex traits patterns may be erroneously explained by homoplasy, that is convergent or parallel evolution. This may be the cases even when coalescent processes that result in incomplete sorting of alleles or traits are accounted for, particularly when the gene flow occurs between relatively distant taxa.

We are indebted to previous work on HRF [12] as the inspiration for our work on G-XRF. HRF is computed per-branch, and we anticipate the development of more granular statistics that apply to local branches, sub-networks, or reticulation nodes within the species network. It is worth noting that as a global metric based on likelihood ratios, G-XRF will reflect the overall risk of introgression. Therefore, a trait pattern with moderate introgression across two clades would have similar risk to that with a high introgression in one clade and a low introgression in the other. As a workaround, researchers may want to compute G-XRF for a particular region or regions of their phylogeny by pruning other taxa. In this way the measure will be more specific and meaningful.

Because we implemented G-XRF using existing multispecies (network) coalescent methods for bi-allelic markers, it does not account for gene duplication and loss or multistate or continuous traits. Previous work on the evolution of quantitative traits within a species tree found that discordance was invariant to the number of loci controlling a trait, a result which may also apply to xenoplasy risk [45]. The framework we presented here is general enough to investigate this and other possibilities, although it requires significant algorithmic improvements. Another useful extension to this framework would be to compute the probabilities where the ancestral state is known, as is the case with Dollo traits where the ancestral state is the presence of a complex trait [46].

We have shown how to visualize the effect on G-XRF when varying up to four parameters in a single analysis (Figs 2 and 4). This will be useful to understand the potential contribution of introgression towards trait patterns when substantial uncertainty is present in one, two, three or four parameters of the model. Greater uncertainty means that a grid search as presented here becomes less feasible, both computationally and in terms of remaining interpretable. Instead, G-XRF could potentially be computed as part of a full Bayesian analysis using MCMC or other algorithms that integrate over the posterior distribution of networks.

Species network inference methods may have trouble identifying instances of reticulate evolution where the introgression probability is very small resulting in a lack of signal, but we do not think this presents a practical problem as such instances necessarily have low xenoplasy risk. The running time for inferring the posterior probability of species networks can be significant; while likelihood calculations for the three-taxon networks took less than one second each, the time complexity of MCMC Bimarkers is O(*sn*^4*l*+4^), where *s* is the number of species, *n* is the number of lineages sampled from all species, and *l* is the level of the network [32, 47]. Increasing the network level is therefore highly deleterious to running time, but this may be overcome using a new, more scalable algorithm with a time complexity of 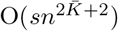, where 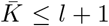 [47]. Another option is using pseudo-likelihood [48], which is much faster to calculate than the full likelihood, though its appropriateness in this domain remains to be studied.

By applying the G-XRF to simulated data, we have demonstrated how the likelihood of particular trait patterns and observed state counts can be meaningfully affected by hybridization and introgression. By applying it to both simulated data and the *Jaltomata* species network, we show how it can be usefully applied by researchers to quantify the risk that particular trait patterns are the product of xenoplasy, instead of or in addition to hemiplasy and homoplasy. Introducing the concept of xenoplasy and a method of estimating the global risk of xenoplasy for binary traits is the first necessary step in developing methods to quantify xenoplasy risk, which we anticipate will flourish given the growing appreciation for the frequency and importance of hybridization and introgression.

## Acknowledgements

This work was funded by National Science Foundation Grants DBI 2030604, CCF 1514177, and CCF 1800723 (to L.N.).

## References

1. Darwin C. On the Origin of Species by Means of Natural Selection, or the Preservation of Favored Races in the Struggle for Life. New York: Modern Library, 1859.

2. Grant PR. Speciation and the adaptive radiation of Darwin’s Finches: the complex diversity of Darwin’s finches may provide a key to the mystery of how intraspecific variation is transformed into interspecific variation. American Scientist 1981;69:653–63.

3. Grant PR and Grant BR. Adaptive radiation of Darwin’s finches: Recent data help explain how this famous group of Galapagos birds evolved, although gaps in our understanding remain. American Scientist 2002;90:130–9.

4. Petren K, Grant P, Grant B, and Keller L. Comparative landscape genetics and the adaptive radiation of Darwin’s finches: the role of peripheral isolation. Molecular Ecology 2005;14:2943–57.

5. Edwards SV. Is a new and general theory of molecular systematics emerging? Evolution: International Journal of Organic Evolution 2009;63:1–19.

6. Nakhleh L. Computational approaches to species phylogeny inference and gene tree reconciliation. Trends in ecology & evolution 2013;28:719–28.

7. Garamszegi LZ. Modern phylogenetic comparative methods and their application in evolutionary biology: concepts and practice. Springer, 2014.

8. Uyeda JC, Zenil-Ferguson R, and Pennell MW. Rethinking phylogenetic comparative methods. Systematic Biology 2018;67:1091–109.

9. Hall BK. Descent with modification: the unity underlying homology and homoplasy as seen through an analysis of development and evolution. Biological Reviews 2003;78:409–33.

10. Tajima F. Evolutionary relationship of DNA sequences in finite populations. Genetics 1983;105:437–60.

11. Avise JC and Robinson TJ. Hemiplasy: A New Term in the Lexicon of Phylogenetics. Systematic Biology 2008;57:503–7.

12. Guerrero RF and Hahn MW. Quantifying the risk of hemiplasy in phylogenetic inference. Proceedings of the National Academy of Sciences 2018;115:12787–92.

13. Liu L, Yu L, and Edwards SV. A maximum pseudo-likelihood approach for estimating species trees under the coalescent model. BMC Evolutionary Biology 2010;10:302.

14. Liu L and Yu L. Estimating species trees from unrooted gene trees. Systematic Biology 2011;60:661–7.

15. Mirarab S, Reaz R, Bayzid MS, Zimmermann T, Swenson MS, and Warnow T. ASTRAL: genome-scale coalescent-based species tree estimation. Bioinformatics 2014;30:i541–i548.

16. Chifman J and Kubatko L. Quartet inference from SNP data under the coalescent model. Bioinformatics 2014;30:3317–24.

17. Ogilvie HA, Bouckaert RR, and Drummond AJ. StarBEAST2 brings faster species tree inference and accurate estimates of substitution rates. Molecular Biology and Evolution 2017;34:2101–14.

18. Flouri T, Jiao X, Rannala B, and Yang Z. Species tree inference with BPP using genomic sequences and the multispecies coalescent. Molecular Biology and Evolution 2018;35:2585–93.

19. Wang Y and Nakhleh LK. Towards an accurate and efficient heuristic for species/gene tree co-estimation. Bioinformatics 2018;34 17:i697–i705.

20. Wang Y, Ogilvie HA, and Nakhleh L. Practical Speedup of Bayesian Inference of Species Phylogenies by Restricting the Space of Gene Trees. Molecular Biology and Evolution 2020;37:1809–18.

21. Hahn MW and Nakhleh L. Irrational exuberance for resolved species trees. Evolution 2016;70:7–17.

22. Maddison WP. Gene Trees in Species Trees. Systematic Biology 1997;46:523–36.

23. Yu Y, Degnan JH, and Nakhleh L. The probability of a gene tree topology within a phylogenetic network with applications to hybridization detection. PLoS Genet 2012;8:e1002660.

24. Yu Y, Dong J, Liu KJ, and Nakhleh L. Maximum likelihood inference of reticulate evolutionary histories. Proceedings of the National Academy of Sciences 2014;111:16448–53.

25. Elworth RL, Ogilvie HA, Zhu J, and Nakhleh L. Advances in computational methods for phylogenetic networks in the presence of hybridization. In: Bioinformatics and Phylogenetics. Springer, 2019:317–60.

26. Karimi N, Grover CE, Gallagher JP, Wendel JF, Ané C, and Baum DA. Reticulate Evolution Helps Explain Apparent Homoplasy in Floral Biology and Pollination in Baobabs (Adansonia; Bombacoideae; Malvaceae). Systematic Biology 2019;69:462–78.

27. Jhwueng DC and O’Meara BC. Trait evolution on phylogenetic networks. bioRxiv 2015:023986.

28. Bastide P, Solís-Lemus C, Kriebel R, William Sparks K, and Ané C. Phylogenetic comparative methods on phylogenetic networks with reticulations. Systematic biology 2018;67:800–20.

29. Hibbins MS, Gibson MJ, and Hahn MW. Determining the probability of hemiplasy in the presence of incomplete lineage sorting and introgression. eLife 2020;9. Ed. by Rokas A and Wittkopp PJ:e63753.

30. Gray GS and Fitch WM. Evolution of antibiotic resistance genes: the DNA sequence of a kanamycin resistance gene from Staphylococcus aureus. Molecular Biology and Evolution 1983;1:57–66.

31. Bryant D, Bouckaert R, Felsenstein J, Rosenberg NA, and RoyChoudhury A. Inferring Species Trees Directly from Biallelic Genetic Markers: Bypassing Gene Trees in a Full Coalescent Analysis. Molecular Biology and Evolution 2012;29:1917–32.

32. Zhu J, Wen D, Yu Y, Meudt HM, and Nakhleh L. Bayesian inference of phylogenetic networks from bi-allelic genetic markers. PLoS computational biology 2018;14:e1005932.

33. Wu M, Kostyun JL, Hahn MW, and Moyle LC. Dissecting the basis of novel trait evolution in a radiation with widespread phylogenetic discordance. Molecular Ecology 2018;27:3301–16.

34. Hudson RR. Generating samples under a Wright–Fisher neutral model of genetic variation. Bioinformatics 2002;18:337–8.

35. Rambaut A and Grass NC. Seq-Gen: an application for the Monte Carlo simulation of DNA sequence evolution along phylogenetic trees. Bioinformatics 1997;13:235–8.

36. Wen D and Nakhleh L. Coestimating reticulate phylogenies and gene trees from multilocus sequence data. Systematic Biology 2017;67:439–57.

37. Zhu J, Yu Y, and Nakhleh L. In the light of deep coalescence: revisiting trees within networks. BMC bioinformatics 2016;17:415.

38. Wen D, Yu Y, Zhu J, and Nakhleh L. Inferring phylogenetic networks using PhyloNet. Systematic Biology 2018;67:735–40.

39. Wen D, Yu Y, Hahn MW, and Nakhleh L. Reticulate evolutionary history and extensive introgression in mosquito species revealed by phylogenetic network analysis. Molecular Ecology 2016;25:2361–72.

40. Wiens JJ. Polymorphism in Systematics and Comparative Biology. Annual Review of Ecology and Systematics 1999;30:327–62.

41. Solís-Lemus C, Yang M, and Ané C. Inconsistency of species tree methods under gene flow. Systematic biology 2016;65:843–51.

42. Miller RJ, Mione T, Phan HL, and Olmstead RG. Color by numbers: Nuclear gene phylogeny of Jaltomata (Solanaceae), sister genus Solanum, supports three clades differing in fruit color. Systematic Botany 2011;36:153–62.

43. Fontaine MC, Pease JB, Steele A, et al. Extensive introgression in a malaria vector species complex revealed by phylogenomics. Science 2015;347.

44. Mallet J, Besansky N, and Hahn MW. How reticulated are species? BioEssays 2016;38:140–9.

45. Mendes FK, Fuentes-González JA, Schraiber JG, and Hahn MW. A multispecies coalescent model for quantitative traits. Elife 2018;7:e36482.

46. Wright AM, Lyons KM, Brandley MC, and Hillis DM. Which came first: The lizard or the egg? Robustness in phylogenetic reconstruction of ancestral states. Journal of Experimental Zoology Part B: Molecular and Developmental Evolution 2015;324:504–16.

47. Rabier CE, Berry V, Glaszmann JC, Pardi F, and Scornavacca C. On the inference of complex phylogenetic networks by Markov Chain Monte-Carlo. bioRxiv 2020.

48. Zhu J and Nakhleh L. Inference of species phylogenies from bi-allelic markers using pseudo-likelihood. Bioinformatics 2018;34:i376–i385.

